# Gut metabolites are more predictive of disease- and cohoused- states than gut bacterial features in a mouse model of polycystic ovary syndrome

**DOI:** 10.1101/2020.10.01.322701

**Authors:** Bryan Ho, Daniel Ryback, Basilin Benson, Pedro J. Torres, Robert A Quinn, Varykina G. Thackray, Scott T. Kelley

## Abstract

Polycystic ovary syndrome (PCOS) impacts ∼10% of reproductive-aged women worldwide. In addition to infertility, women with PCOS suffer from metabolic dysregulation which increases their risk of developing type 2 diabetes, cardiovascular disease and non-alcoholic fatty liver disease. Studies have shown differences in the gut microbiome of women with PCOS compared to controls, a pattern replicated in mouse models. Recently, using a letrozole-induced mouse model of PCOS, we demonstrated that cohousing was protective against development of metabolic and reproductive phenotypes and showed via 16S amplicon sequencing that this protection correlated with time-dependent shifts in gut bacteria. Here, we applied untargeted metabolomics and shotgun metagenomics approaches to further analyze the longitudinal samples from the cohousing experiment. Analysis of beta diversity found that untargeted metabolites had the strongest correlation to both disease and cohoused states and that shifts in metabolite diversity were detected prior to shifts in bacterial diversity. In addition, log2-fold analyses found numerous metabolite features, particularly bile acids (BA), to be highly differentiated between placebo (P) and letrozole (LET), as well as cohoused LET versus LET. Our results indicate that changes in gut metabolites, particularly BAs, are associated with a PCOS-like phenotype in the LET mouse model as well as the protective effect of cohousing. Our results also suggest that transfer of metabolites via coprophagy occurs rapidly and may precipitate changes in bacterial diversity. This study joins a growing body of research highlighting changes in primary and secondary bile acids that may provide a link between host metabolism and gut microbes relevant to the pathology of PCOS.

**IMPORTANCE:** Using a combination of untargeted metabolomics and metagenomics, we performed a comparative longitudinal analysis of the feces collected in a cohousing study with a PCOS mouse model. Our results showed that gut metabolite composition experienced earlier and more pronounced differentiation in both the disease model and cohoused mice compared with the microbial composition. Notably, statistical and machine learning approaches identified shifts in the relative abundance of primary and secondary BA, which have been implicated as modifiers of gut microbial growth and diversity. Network correlation analysis showed strong associations between particular BA and bacterial species, particularly members of *Lactobacillus*, and that these correlations were time and treatment dependent. Our results provide novel insights into host/microbe relationships related to hyperandrogenism in females and indicate that focused research into small molecule control of gut microbial diversity and host physiology may provide new therapeutic options for the treatment of PCOS.

## INTRODUCTION

Polycystic ovary syndrome (PCOS), a common reproductive endocrine disorder, is estimated to affect ∼5-15% of reproductive age women worldwide (1). PCOS is the most prevalent cause of anovulatory infertility and women with this disorder have a higher risk of pregnancy-related complications (2). The diagnosis of PCOS is based on the 2003 Rotterdam criteria, which requires two out of three criteria: hyperandrogenism, oligomenorrhea or amenorrhea, and polycystic ovaries (3). Although the precise etiology of PCOS is unknown, genetic and twin studies indicate that PCOS is a polygenic heritable disorder that is influenced by environmental factors including exposure to excess maternal androgens during fetal development (4–6). The onset of PCOS often occurs during the early reproductive years, indicating that puberty may be a critical period in the development of PCOS (7).

In addition to its effects on reproductive health, PCOS increases the risk of developing metabolic diseases such as type 2 diabetes, hypertension and non-alcoholic fatty liver disease (NAFLD) (8, 9). Metabolic dysregulation manifests predominantly in women with PCOS that have hyperandrogenism and is independent of body mass index (10, 11). Alongside metabolic dysregulation, PCOS is also associated with changes in the gut microbiome (12, 13). The gut microbiome comprises a complex community of microorganisms that are important for host physiology including immunity, metabolism, and neurology (14). Gut microbes play a critical role in the fermentation of dietary fibers, synthesis of vitamins such as B12, modification of bile acids, neurotransmitters and hormones, and production of short chain fatty acids that regulate energy homeostasis (14). Dysbiosis of the gut microbiome has been correlated with multiple metabolic disorders, including obesity, type 2 diabetes, and NAFLD (15). With regards to PCOS, studies have shown that gut bacterial species richness is lower and that the relative abundance of specific bacterial taxa is altered in adolescent girls or women with PCOS compared to women without the disorder (12, 13). Furthermore, studies have demonstrated a strong correlation between gut microbial diversity or the abundance of specific gut bacterial taxa and hyperandrogenism, indicating that testosterone may modulate the composition of the gut microbiome in women (16).

Investigations of the gut microbiome in a letrozole-induced PCOS mouse model have also indicated a strong relationship between hyperandrogenism and shifts in the alpha diversity and composition of the gut microbiome (17, 18). This mouse model utilizes letrozole, a non-steroidal aromatase inhibitor, to limit the conversion of testosterone to estrogen, resulting in increased testosterone and decreased estrogen levels. This model recapitulates both reproductive and metabolic hallmarks of PCOS including elevated luteinizing hormone (LH) levels, oligo-or anovulation, polycystic ovaries, weight gain, abdominal adiposity, dysglycemia, hyperinsulinemia, and insulin resistance (17–19). The importance of hyperandrogenism in this activational model was further demonstrated in a study that discontinued letrozole treatment in mice and demonstrated a recovery in reproductive, metabolic, and gut microbial phenotypes (20). A letrozole-induced PCOS rat model were also reported to have changes in the gut microbiome (21, 22).

Although correlative evidence from both human and rodent model studies indicates that there is an association between PCOS and the gut microbiome, a direct role of the gut microbiome in generating or exacerbating metabolic dysregulation in this disease state has yet to be established. Indeed, despite the many studies that have identified correlations between metabolic disorders such as obesity, type 2 diabetes, and NAFLD and shifts in the gut microbiome, very few have demonstrated a direct effect of gut microbes in these disorders (23). One method for establishing a causal link between the gut microbiome and host is via fecal microbiota transplant (FMT) into germ-free mice. For example, Ridaura et al. transplanted stool samples from lean and obese human donors into germ-free mice and found that the mice developed the donors’ metabolic phenotype (24). More recently, Qi et al. performed a FMT of stool from women with PCOS versus controls into antibiotic-depleted mice and showed that the FMT with PCOS stool was sufficient to result in a PCOS-like phenotype that included increased LH, acyclicity, polycystic ovaries, and insulin resistance (25). Since rodents are coprophagic, another approach is to perform a cohousing study. This method of horizontal transmission has been shown to result in exchange of microbiota between caged individuals (26). Cohousing with healthy mice was reported to be protective against developing obesity and maternal high-fat diet induced metabolic dysregulation (24, 27). Altogether, these studies suggest that the gut microbiome may play a causative role in the development of metabolic disorders including PCOS.

Recently, we performed a cohousing study using the letrozole mouse model to test whether exposure to a healthy gut microbiome was protective against developing PCOS metabolic or reproductive phenotypes (19). This study consisted of four groups: placebo mice housed together (P), letrozole mice housed together (LET), placebo mice cohoused with letrozole mice (P^ch^), and letrozole mice cohoused with placebo mice (LET^ch^). This study demonstrated that cohousing with healthy mice improved both reproductive and metabolic phenotypes associated with letrozole treatment while placebo mice cohoused with letrozole mice did not exhibit any PCOS complications (19). Interestingly, gut microbial 16S rRNA sequencing analysis showed that the gut microbiome of LET^ch^ mice did not resemble the gut microbiome of placebo mice but instead was more similar to P^ch^ (19). These results suggested that cohousing resulted in an exchange of gut microbes and that transfer of the gut microbiome from a healthy mouse to a letrozole-treated mouse was sufficient to provide protection from developing both metabolic and reproductive phenotypes of PCOS despite the lack of similarity between the placebo and LET^ch^ mice.

In the present study, we applied multiple ‘omics approaches to further explore the longitudinal relationship of the gut microbiome samples collected in the LET-induced PCOS mouse model cohousing study (Torres ref). Although little is known about the relationship between gut metabolites and PCOS, a recent study by Qi *et al*., 2019 showed that the secondary bile acids, glycodeoxycholic acid and tauroursodeoxycholic acid were present in lower abundance in a cohort of women with PCOS relative to women without this disorder, and that supplementation with these bile acids was protective against developing a PCOS-like phenotype in mice (Qi et al 2019). Given these results, we hypothesized that gut metabolites might provide greater explanatory power for longitudinal patterns in the data than gut microbes and that the combination of multiple ‘omics datasets might strengthen the time-dependent patterns previously observed with the 16S data. Specifically, we applied untargeted mass spectrometry analyses to identify the diversity of small molecules, including bile acids, in the fecal samples, and shotgun metagenomic sequencing to improve species and strain level identification of bacterial species.

## RESULTS

### Metabolites clustered by treatment in a time-dependent manner

Add sentence about how much features identified (known versus unknown). A canonical analysis of principal coordinates (CAP) analysis of untargeted metabolomic data generated from weekly fecal samples from the cohousing study found the degree of clustering associated with treatment differed substantially over the course of the study (Fig. 1). No treatment-associated clustering was observed based on sample metabolite composition prior to pellet implantation (week 0) (Fig. 1A). By week 1, we observed differentiation among samples from different treatments (Fig. 1B; p=0.002, R^2^=0.161), with virtually no overlap among samples from the four treatment groups. The greatest degree of clustering occurred among the week 2 samples (Fig. 1C; p=0.001, R^2^=0.427). At week 2, P and LET samples showed distinct separation from each other and from the cohousing samples (P^ch^ and LET^ch^) which clustered together. The amount of variation explained by the first two principal components was also highest at week 2 (Fig. 1C). We continued to observe significant associations between treatment and metabolites at week 3, but this was not observed at weeks 4 and 5 of the study (Figs. 1D-F).

**FIG 1.**
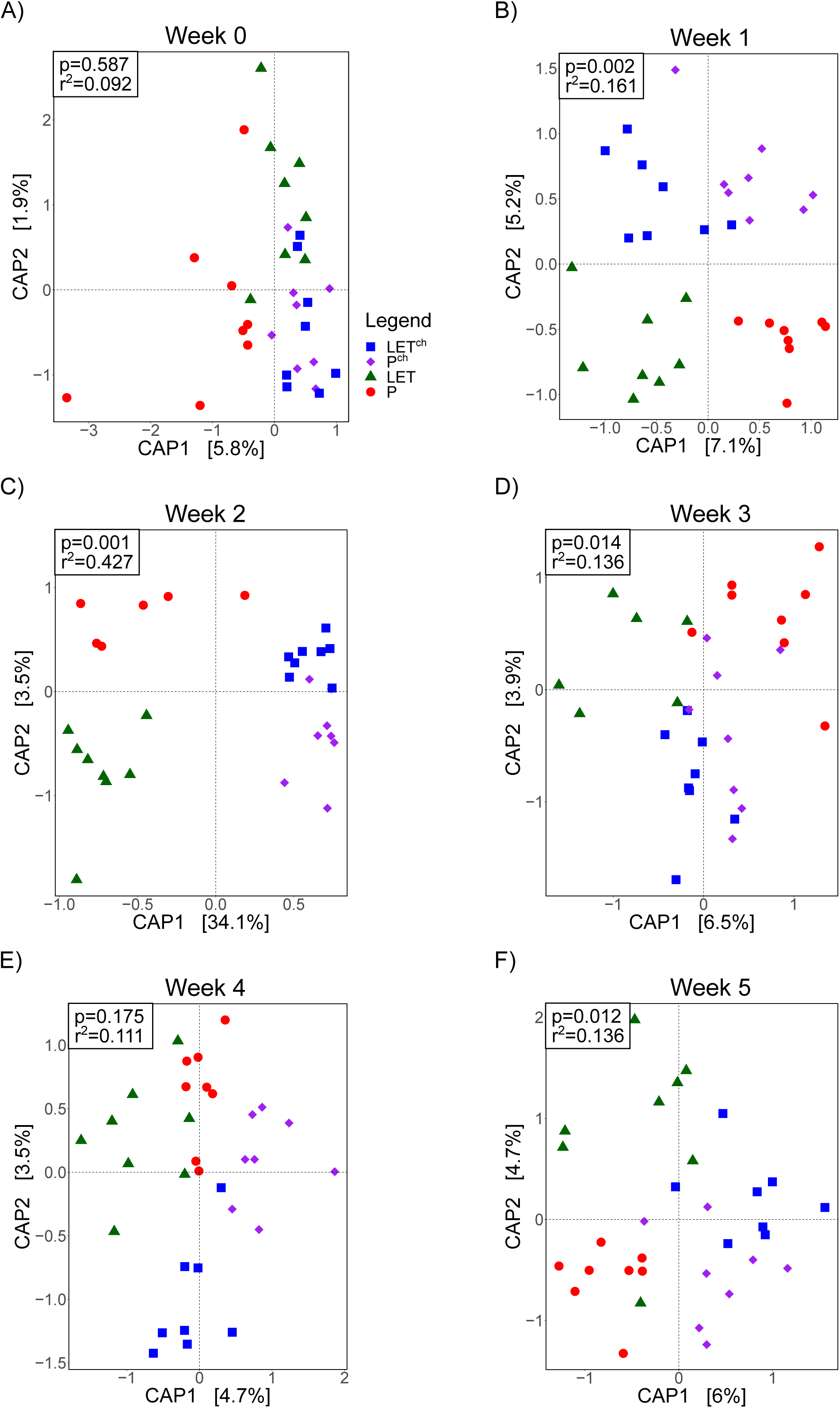
Cohousing letrozole-treated mice with placebo-treated mice influenced the overall composition of gut microbial metabolites over time. The cohousing study included four groups of mice (n=8/group): placebo (P), letrozole (LET), placebo mice cohoused with letrozole (P^ch^) and letrozole mice cohoused with placebo (LET^ch^). (A-F) Constrained canonical analysis of principal (CAP) coordinates (Bray-Curtis distances) of fecal metabolites from the four treatment groups for weeks 0-5 post-treatment. Shown are the CAP1 and CAP2 coordinates, representing the two coordinates that captured the greatest amount of variation (percentage of variation is shown in brackets). Results of Permutational ANOVA (PERMANOVA) analysis of the Bray-Curtis distances are shown for each time point.

### Comparisons of beta diversity analyses across multiomics datasets found strongest association between metabolites and treatment

The metabolite CAP analysis was compared with metagenomic and 16S rRNA CAP analyses of the gut microbiome at weeks 2 and 5, the time points for which we had all three data sets. At week 2, all three datasets showed clear differentiation among the samples from the four treatment groups (Fig. 2). The goodness of fit (R^2^) between data and treatment was highest for the metabolites (Fig. 2A; p=0.001, R^2^=0.427) followed by the 16S (Fig 2C; p=0.027, R^2^=0.207) and metagenomic (Fig 2E; p=0.036, R^2^=0.193) data. The proportion of variation explained by the first two principal components at week 2 was also the highest with the metabolite data compared to the metagenome and 16S data. At week 5, the metabolites showed significant levels of differentiation among treatments (Fig. 2B; p=0.012, R^2^=0.136), while we found no significant separation based on the 16S (Fig. 2D; p=0.320, R^2^=0.171) and metagenomic (Fig. 2F; p=0.072, R^2^=0.153) data. For all three datasets, both the goodness of fit (R^2^ values) and the amount of variation explained by the first two principal components was lower at week 5 than week 2.

**FIG 2.**
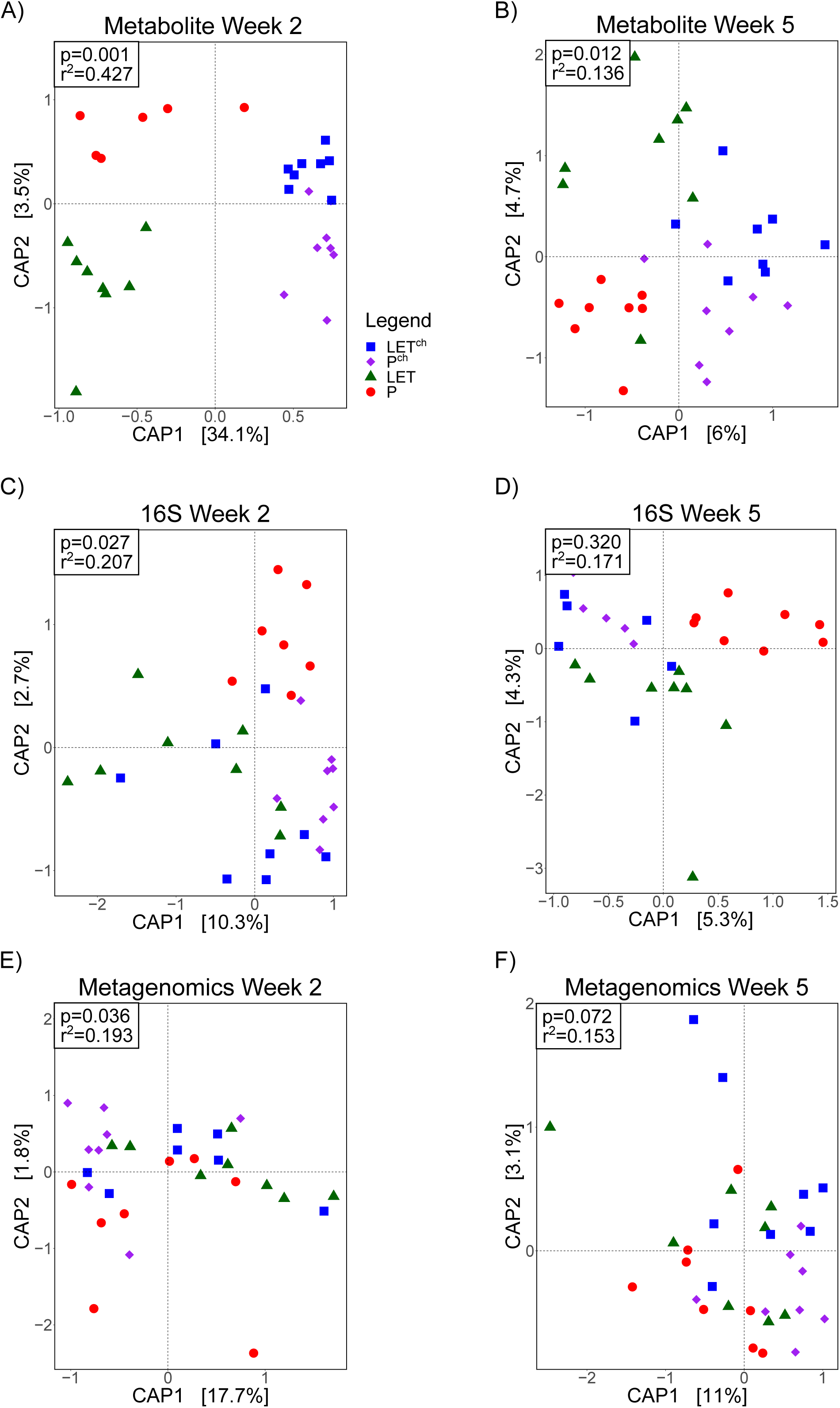
Cohousing letrozole with placebo mice resulted in greater differentiation among treatment groups in the overall composition of both gut microbes and metabolites at 2 weeks compared with 5 weeks. The cohousing study included four groups of mice: placebo (P), letrozole (LET), placebo mice cohoused with letrozole (P^ch^) and letrozole mice cohoused with placebo (LET^ch^). Canonical analysis of principal (CAP) coordinates of Bray-Curtis dissimilarity among the four treatment groups, Week 2 (A, C, E) and Week 5 (B, D, F). Shown are the CAP1 and CAP2 coordinates, representing the two coordinates that captured the greatest amount of variation (percentage of variation is shown in brackets). (A-B) metabolites (C-D) bacterial 16S rRNA gene sequences (16S) (E-F) bacterial whole genome sequencing (WGS).

### Combining multiomics datasets did not improve goodness-of-fit in beta diversity analyses

CAP analyses were performed after combining the metabolomics data with the 16S and metagenomic datasets at weeks 2 and 5, respectively. At week 2, the combined metabolite and 16S data showed clear differentiation between treatment groups (Fig. 3A; p=0.001, R^2^=0.425) but was not different at week 5 (Fig. 3B; p=0.060, R^2^=0.124). For the combined metabolite and metagenomic dataset, treatment groups were significantly different at both week 2 (Fig. 3C: p=0.001, R^2^=0.398) and week 5 (Fig. 3D; p=0.004, R^2^=0.135). The amount of variation explained by the first two principal components was higher at week 2 for both combined datasets. The goodness-of-fit and amount of variation explained by the principal components at week 2 were not greater than determined independently for the metabolite dataset (Fig 2A), indicating that the metabolite dataset in this multiomics analysis was the main contributor to the observed treatment differentiation.

**FIG 3.**
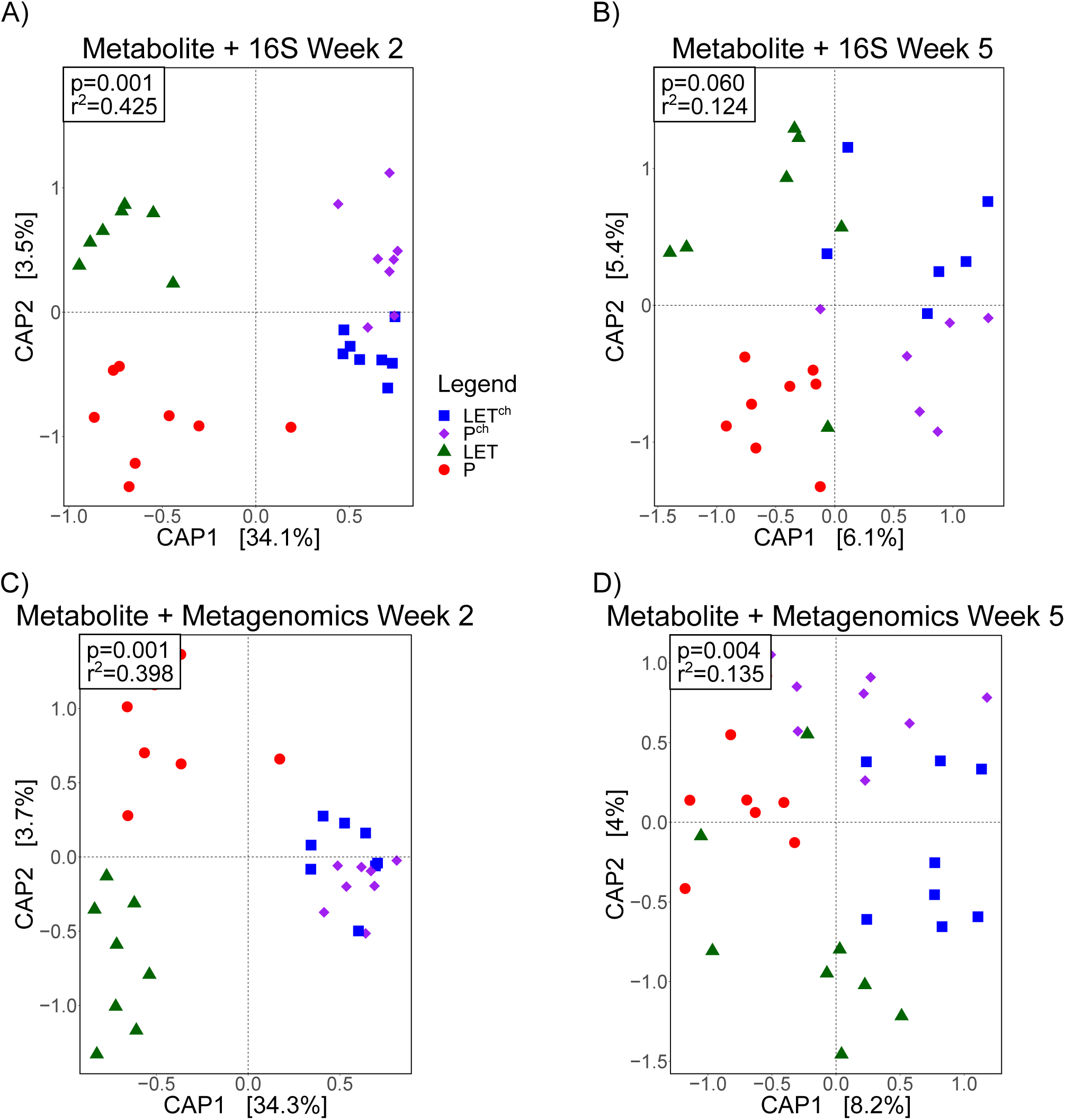
Combination of metabolomic and metagenomic data did not improve the fit of overall microbial compositional data compared to metabolomic dataset alone. The cohousing study included four groups of mice: placebo (P), letrozole (LET), placebo mice cohoused with letrozole (P^ch^) and letrozole mice cohoused with placebo (LET^ch^). Canonical analysis of principal (CAP) coordinates of Bray-Curtis dissimilarity among the four treatment groups, Week 2 (A, C) and Week 5 (B, D). Shown are the CAP1 and CAP2 coordinates, representing the two coordinates that captured the greatest amount of variation (percentage of variation is shown in brackets). (A-B), metabolites combined with bacterial 16S rRNA gene sequences (16S), (C-D), metabolites combined with bacterial whole genome sequencing (WGS).

### Log_2_ fold analysis identified numerous differential metabolite abundances between treatment groups

To identify specific features that contributed to the difference in treatment groups, we calculated log_2_ fold ratios for the relative abundances of primary and secondary BAs and other identifiable metabolites (Fig. 4). For the BAs, the number of differential BAs was greater between LET^ch^ and LET than between P and LET, particularly at week 2 (Fig. 4 A-D). We found a similar, though more pronounced pattern with the other identified metabolites (Fig. 4 E-H). Almost twice as many identifiable metabolites were differentially abundant in LET^ch^ /LET compared to P/LET at week 2 (Fig. 4E, F) but this pattern was not present at week 5 (Fig. 4G, H).

**FIG 4.**
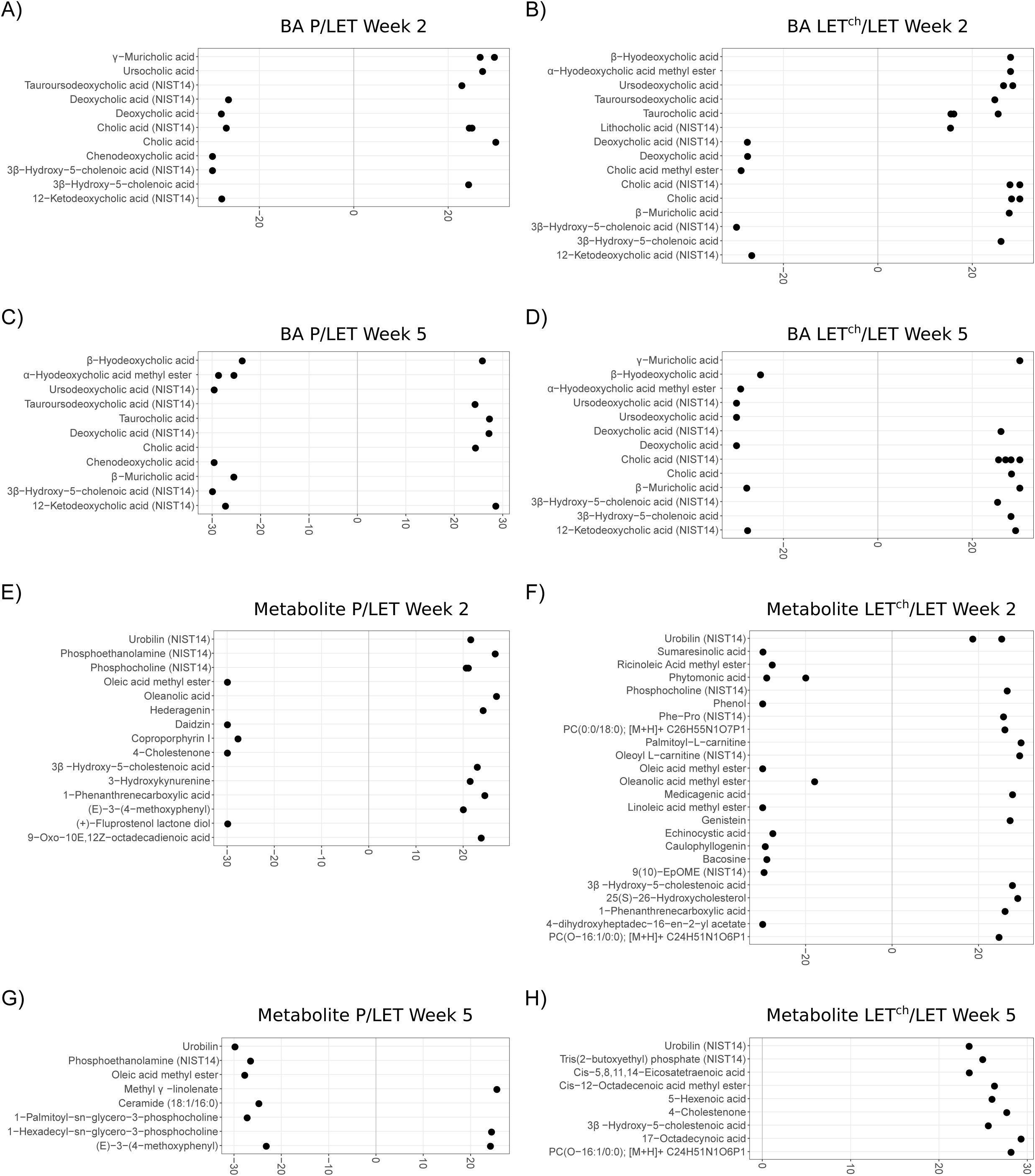
Primary and secondary bile acid relative abundance was altered in a pairwise comparison of P relative to LET or LET^ch^ relative to LET. The cohousing study included four groups of mice: placebo (P), letrozole (LET), placebo mice cohoused with letrozole (P^ch^) and letrozole mice cohoused with placebo (LET^ch^). Results from the DeSeq2 analysis were expressed as log2 fold change for P/LET (A, C, E, G) and LET^ch^/LET (B, D, F, H) with (A-B) bile acids (BA) Week 2, (C-D) BA Week 5, (E-F) other identified metabolites Week 2, and (G-H) other identified metabolites Week 5. The ten BA or metabolites with the greatest magnitude of log2 fold change were shown for each comparison. Positive log2 fold changes represent metabolites increased in P or LET^ch^ relative to LET, while negative changes represent metabolites increased in LET relative to P or LET^ch^.

### Log_2_ fold change values smaller with bacterial relative abundances compared to metabolites

Figure 5 shows the results of log_2_ fold analyses based on relative bacterial abundances estimated via 16S rRNA gene sequences (Fig. 5 A-D) and metagenomes (Fig. 5 E-H). The plots include the ten taxa with the greatest fold change from each comparison. The fold change values ranged from −6 to +6 with the 16S OTU data, and −3 to +2 with the metagenomic data, compared with a fold change range of −30 to 30 with the metabolites (Fig. 4). In the 16S data, comparisons of week 2 and 5 found 6 of the top 10 taxa with the highest fold change values were the same for both P/LET and LET^ch^/LET comparisons. At week 2, P/LET and LET^ch^/LET shared 6 of the same OTUs, and all but Ruminococcus had a similar positive or negative change in ratio. For the metagenomic data, we identified 4 bacterial species shared between week 2 and 5 for P/LET and 2 species shared between weeks 2 and 5 in the LET^ch^/LET comparisons. Only 2 species, *Akkermansia munciniphilia* and *Pseudobutyrivibiro ruminis* were in common at week 2 between the P/LET and LET^ch^/LET comparisons.

**FIG 5.**
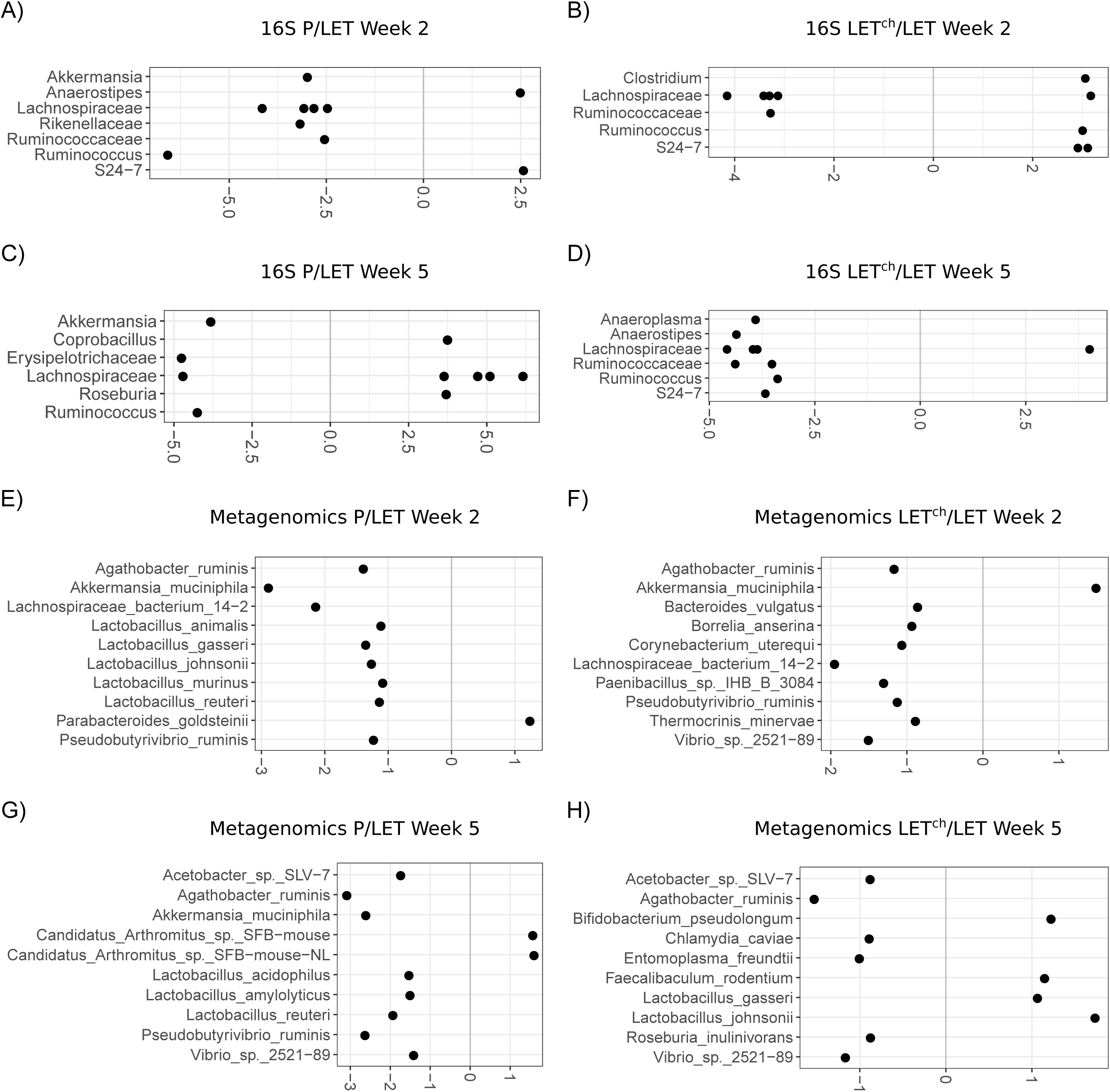
Bacterial relative abundance was altered in a pairwise comparison of P relative to LET or LET^ch^ relative to LET. The cohousing study included four groups of mice: placebo (P), letrozole (LET), placebo mice cohoused with letrozole (P^ch^) and letrozole mice cohoused with placebo (LET^ch^). Results from the DeSeq2 analysis were expressed as log2 fold change for P/LET (A, C, E, G) and LET^ch^/LET (B, D, F, H) with (A-B) 16S rRNA bacterial gene sequences (16S) Week 2, (C-D) 16S Week 5, (E-F) bacterial whole genome sequencing (WGS) Week 2, and (G-H) WGS Week 5. The ten bacteria (family or genus level for 16S; species level for WGS) with the greatest magnitude of log2 fold change were shown for each comparison. Positive log2 fold changes represent bacteria increased in P or LET^ch^ relative to LET while negative changes represent bacteria increased in LET relative to P or LET^ch^.

### Multiomic data combinations improved random forest classification accuracy

Table 1 shows the results of random forest analyses classification of P vs. LET and LET^ch^ vs. LET comparisons at week 2 and 5 for independent and combined datasets. Generally speaking, the week 2 datasets were better able to classify treatments than week 5, and classification accuracy was highest with LET^ch^ vs. LET. Among independent datasets, 16S dataset had the highest classification accuracy, while the combination of BA and metagenomic data resulted in the best overall accuracy for both P vs. LET and LET^ch^ vs. LET comparisons. An analysis of features that contributed the most to the accuracy of the combined multiomics random forest analyses is shown in Figure 6. Specifically, the 10 features with the highest Gini importance scores are shown; their removal from the dataset has the greatest effect on the ability to classify between the treatment conditions. In the week 2 samples, BAs were the majority of the top 10 features with the highest Gini importance. In the LET^ch^ vs. LET comparisons, all of the most importance classification features were BAs when they were combined with 16S data (Fig. 6B), while 8 of the 10 most important features were BAs when they were combined with metagenomic data (Fig. 6F). At week 2, 6 out of the 10 more important features were BAs in the P vs. LET comparisons at week 2 for both the BA + 16S combination (Fig. 6A) and the BA + metagenome combination (Fig. 6E). In the week 5 samples, the case was reversed: with one exception (Fig. 6D, BA +16S LET^ch^ vs. LET) the bacterial taxa dominated the top 10 importance features.

**TABLE 1.** Results of Random Forest analyses with single and combined multiomics datasets.

| Comparisons | Accuracy<sup>a</sup> | OOB Score<sup>b</sup> | Mean Accuracy<sup>c</sup> | AUC<sup>d</sup> |
| --- | --- | --- | --- | --- |
| **Week 2** |  |  |  |  |
| P vs LET (BA) | 16.67% | 50.00% | 40.00% | 0.00% |
| P vs LET (16S) | 83.33% | 20.00% | 70.00% | 66.66% |
| P vs LET (WGS) | 66.67% | 80.00% | 70.00% | 100.00% |
| LET vs LETch (BA) | 66.67% | 80.00% | 80.00% | 100.00% |
| LET vs LETch (16S) | 100.00% | 93.75% | 90.00% | 100.00% |
| LET vs LETch (WSG) | 66.67% | 40.00% | 40.00% | 66.67% |
| P vs LET (BA+16S) | 50.00% | 50.00% | 60.00% | 66.66% |
| P vs LET (BA+WGS) | 100% | 60.00% | 60.00% | 100.00% |
| LET vs LETch (BA+16S) | 100.00% | 100.00% | 90.00% | 100.00% |
| LET vs LETch (BA+WGS) | 100% | 90.00% | 80.00% | 100.00% |
| **Week 5** |  |  |  |  |
| P vs LET (BA) | 50.00% | 80.00% | 80.00% | 66.67% |
| P vs LET (16S) | 50.00% | 40.00% | 60.00% | 55.55% |
| P vs LET (WGS) | 66.67% | 90.00% | 70.00% | 55.56% |
| LET vs LETch (BA) | 66.67% | 80.00% | 70.00% | 44.45% |
| LET vs LETch (16S) | 40.00% | 30.00% | 73.33% | 16.66% |
| LET vs LETch (WSG) | 66.67% | 60.00% | 70.00% | 88.89% |
| P vs LET (BA+16S) | 50.00% | 30.00% | 60.00% | 66.66% |
| P vs LET (BA+WGS) | 83.30% | 60.00% | 90.00% | 78.00% |
| LET vs LETch (BA+16S) | 40.00% | 77.77% | 73.33% | 83.33% |
| LET vs LETch (BA+WGS) | 83.30% | 30.00% | 50.00% | 88.90% |

P=Placebo, LET=Letrozole, LETch= LET cohoused with Placebo.
BA=Bile Acids; 16S=16S rRNA gene sequences; WGS=Whole Genome sequencing data.
<sup>a</sup>RF Accuracy = Percentage of correctly classified samples.
<sup>b</sup>Out of Bag Score = Percentage of correctly classified samples based on unselected samples from the bootstrap sampling method.
<sup>c</sup>Mean Accuracy = Average accuracy of a 5-fold cross validation.
<sup>d</sup>Area under the curve = Measurement of the performance of a binary classifier.

**FIG 6.**
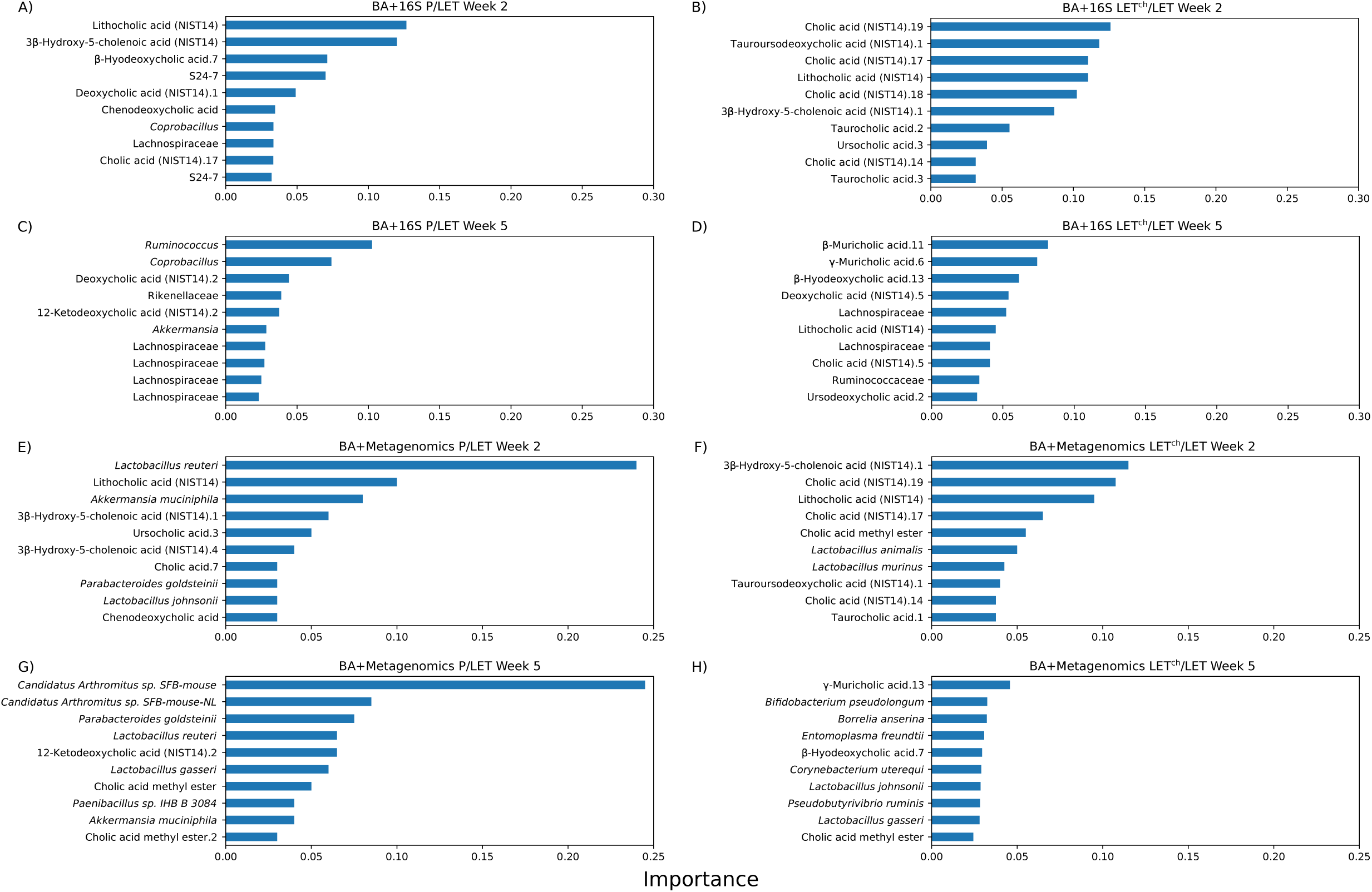
Top ten bile acid and bacterial features that classify P versus LET or LET^ch^ versus LET in Random Forest analysis. The cohousing study included four groups of mice: placebo (P), letrozole (LET), placebo mice cohoused with letrozole (P^ch^) and letrozole mice cohoused with placebo (LET^ch^). The graphs show Gini importance scores which indicate the relative importance a particular feature [bile acid (BA) or bacteria] in the classification result. Results are shown for P/LET (A, C, E, G) and LET^ch^/LET (B, D, F, H) with (A-B) BA+bacterial 16S rRNA gene sequencing (16S) Week 2, (C-D) BA+16S Week 5, (E-F) BA+whole genome sequencing (WGS) Week 2, and (G-H) BA+WGS Week 5.

### Analyses revealed time and treatment specific patterns of correlations between bacteria and bile acids

The heatmaps in Figure 7 illustrate the results of correlations between BAs and bacterial species identified in the metagenomic data. For all the treatment groups (P, LET and LET^ch^), we observed more than twice the number of strong correlations (p ≥ |0.8|; dark red or blue squares) in samples collected at week 2 than in those collected at week 5. Moreover, none of the strongest correlations detected between BAs and bacterial species at week 2 were detectable at week 5. For example, in the week 2 P samples, one cholic acid metabolite was positively correlated with 6 different bacterial species. However, this same metabolite was weakly or even negative correlated with the same 6 species at week 5. Similar patterns could be observed in the LET (e.g., γ-Muricholic acid) and LET^ch^ (e.g., Cholic acid.7) treatment groups. The patterns of correlations between bacteria and BAs also differed considerable among treatment groups. Of the ten strongest correlations between BAs and bacterial species in the P samples at week 2, 8 were not detectable in the LET samples. Similarly, none of the ten strongest correlations in LET^ch^ at week 2 were correlated in LET at the same time point.

**FIG 7.**
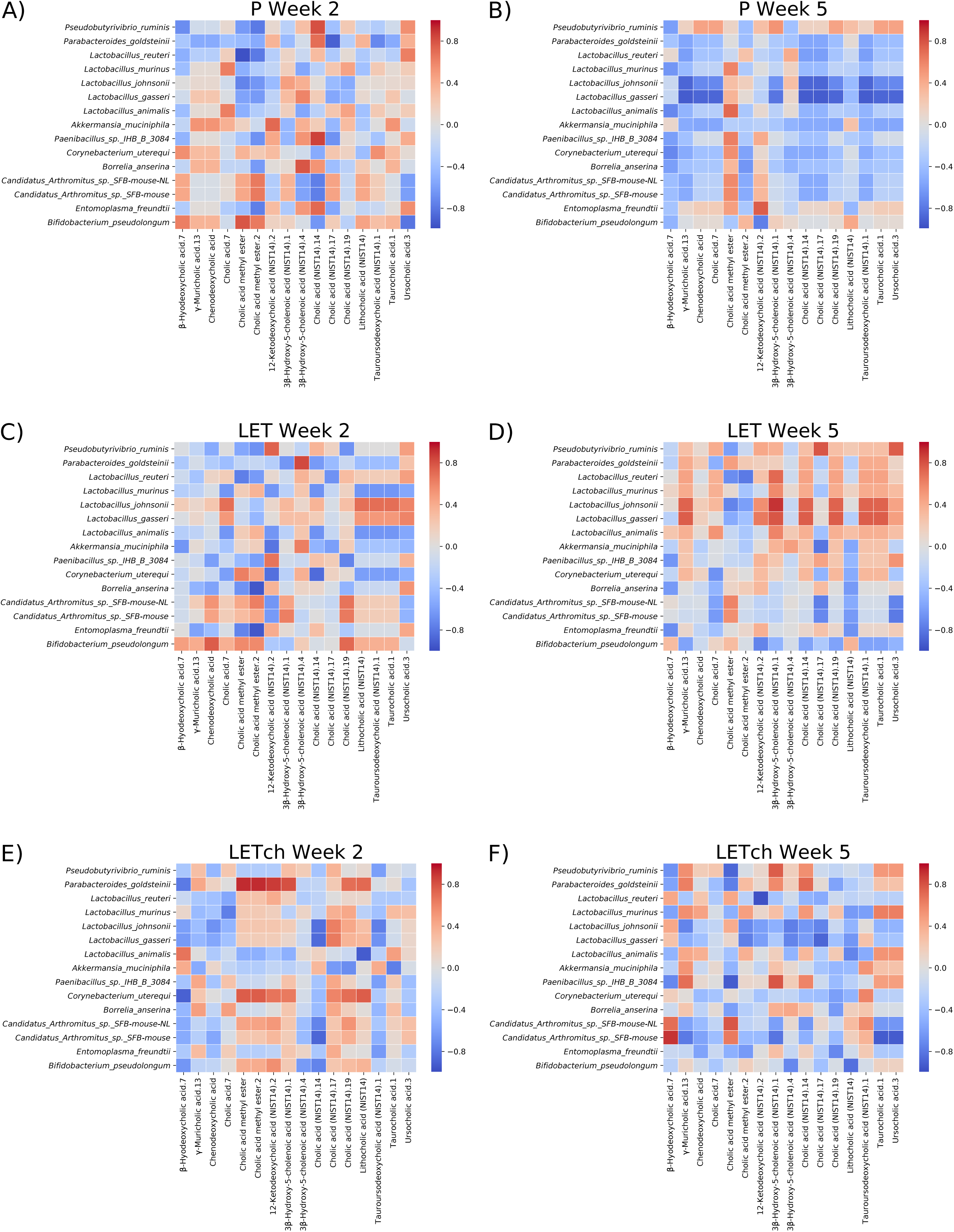
Heatmaps of Spearman-rank correlation values between clr-transformed bile acid and bacterial abundances. Red indicates positive correlation values, while blue indicates negative correlation values. The bile acid and bacterial features chosen for analysis were selected based on DeSeq2 analyses. The cohousing study included four groups of mice: placebo (P), letrozole (LET), placebo mice cohoused with letrozole (P^ch^) and letrozole mice cohoused with placebo (LET^ch^). (A, B) P, (C,D) LET and (E, F) LET^ch^ Weeks 2 or 5.

## DISCUSSION

Multiomics methods improved the fit of gut microbiome samples to the experimental treatment conditions, increased the resolution of the time-dependent changes in the gut microbiome, and identified potential targets for future mechanistic studies. The untargeted metabolomic analysis proved especially useful for distinguishing treatment groups. At time 0, of the longitudinal sampling, there were no detectable differences among treatment groups (Fig. 1A), but by week 1, the untargeted metabolomic data indicated clear patterns of differentiation (Fig. 1B). By week 2, all three data sets showed a pattern of differentiation, but the metabolite data was the best fit to the treatment conditions (Fig. 1C). Analysis of the untargeted metabolites from the cohousing samples also revealed a pattern of early divergence of both P^ch^ and LET^ch^ from the P and LET microbiome samples, which were also distinct from one another. Furthermore, effects of cohousing on the LET^ch^ gut microbiome did not result in a “placebo-like” state; rather cohousing resulted in a third state, distinct from both P and LET conditions, indicating that the gut microbiome does not have to “return” to the healthy state in order to have protective effects.

The time-dependent nature of the shift in the gut microbiome due to both LET treatment and cohousing was apparent in all the -omics datasets. Untargeted metabolomics, 16S and shotgun metagenomics showed stronger separation at week 2 samples than week 5, with the metabolite data being the best fit to treatment at both time points (Fig. 2). These results further confirm the importance of longitudinal sampling. Had we waited until the end of the experiment to collect fecal samples, or sampled at only a single time point, our interpretation of the results would have been affected.

Overall, three clear patterns emerged from the longitudinal multiomics analyses: (1) The shift in overall gut metabolite diversity was detectable prior to the shift in bacterial diversity (16S or metagenomics); (2) A detectable shift in gut metabolites happened by the first week of the study; and (3) Both P^ch^ and LET^ch^ were distinct from the P and LET samples, respectively and were also more similar to one another than either were to the P and LET samples. The early community-wide shift in metabolite diversity in the LET samples suggests that these changes began even earlier, perhaps immediately after letrozole treatment and the resultant increase in testosterone levels. Further, our results suggest that metabolites transferred quickly between P^ch^ and LET^ch^ mice, likely via coprophagy. Given these patterns, we hypothesize that host-related changes led to changes in gut microbial diversity, though future experimentation will be needed to understand the timing and nature of these changes.

While little is currently known about the role specific bile acids have in modulating host/microbe interactions, it is well known that bile acids can affect the growth of gut bacteria and that they are chemically modified by gut bacteria. Bile acids have been reported to promote the growth of bile-acid metabolizing bacteria and have strong antibiotic properties (28) and favor resistant bacteria such as *Lactobacillus* and *Bifidobacterium* (29). Numerous gut bacteria including members of the *Lactobacillus, Bifidobacterium* and *Clostridium* also express bile-salt hydrolase (BSH) enzymes, enabling them to deconjugate BAs (30). In addition, *Clostridium* species can metabolize secondary bile acids (30). Interestingly, our log_2_ fold analysis found that many species within these bacteria genera (e.g., *Lactobacillus, Bifidobacterium, Akkermansia*, and *Clostridium*) were differentially abundance between P and LET as well as LET and LET^ch^, particularly at week 2 (Fig. 5). A recent study by Tian *et al*., 2019 measured the effects of BAs on the growth and survival of four common gut bacteria and found that different BAs had bacterial-specific effects (31). Interestingly, the BAs they tested in the study, lithocholic acid (LCA), deoxycholic acid (DCA), taurocholic acid (TCA), and tauroursodeoxycholic acid (TUDCA), were identified as differentially abundant in log_2_ fold comparisons between P/LET and LET^ch^/LET (Fig. 4) and had high Gini importance scores in the random forest analyses, particular at week 2 (Fig. 6A, B). In addition, our study identified strong correlations between 3 of these BAs (LCA, TCA and TUDCA) and various gut bacteria in our study (Fig. 7). Furthermore, the magnitude of these correlations varied considerably among treatment groups. For example, in the LET samples at week 2 LCA, TCA and TUCA were highly positively correlated with *Lactobacillus johnsonii* and *L. gasseri* and strongly negatively correlated with *L. murinus* and *L. animalis* (Fig. 7C). At the same time point there were no correlation detected between these three BAs and these *Lactobacillus* species in P samples (Fig. 7A) and only strong negative correlations between TCUA and *L. johnsonii* and *L. gasseri* in LET^ch^ (Fig. 7E). Interestingly, a recent study of the effects of BAs on the neonatal microbiome found that administration of specific BAs to mice via oral gavage had a particularly strong effects on members of the *Lactobacillus*, and that this effect was highly species dependent (32) *L. johnsonii*, which carries BSH enzymes of different specificities (33), was strongly affected by the administration of multiple different BAs, while the BSH-negative *L. murinus* was unaffected. Two other striking patterns were the strong positive correlations in the week 2 LET^ch^ samples between the same four bile acids and *Parabacteroides goldsteinii* and, to a lesser extent, *Corynebacterium uterequi* (Fig. 7E), correlations not observed in either LET or P samples. Species of *Parabacteroides* have been suggested as potential therapeutics for obesity-related disorders and have been shown to metabolize BAs (34).

Overall, our study joins a small but growing body of literature indicating that changes in primary and secondary bile acids may be associated with PCOS. Somewhat surprisingly given the importance of bile acids in host physiology, much remains to be discovered concerning the mediation of bile acid signaling by host-microbe interactions. With regards to PCOS, this will necessitate mechanistic studies in animal models focused on the role of sex steroids in regulating BA production in the liver, BA metabolism by gut microbes and bile acid signaling in the enterohepatic system.

## MATERIALS AND METHODS

### Letrozole-induced PCOS mouse model

Details on the mouse model and cohousing experimental design were described previously (19). Briefly, C57Bl/6NHsd female mice (Envigo) were housed in a vivarium under specific pathogen-free conditions with *ad libitum* access to water and food. Placebo or 3 mg letrozole pellets (50 µg/d; Innovative Research of America) were implanted subcutaneously into four-week-old mice for five weeks. Throughout the experiment, mice were housed two per cage in three separate arrangements (n=8 mice/group, 32 total): two placebo mice (P), two letrozole mice (L), or cohoused one placebo (P^ch^) and one letrozole (LET^ch^) mouse. All animal procedures in the experiment were approved by the University of California, San Diego Institutional Animal Care and Use Committee (Protocol: S14011).

### Fecal sample collection, DNA isolation and 16S rRNA gene sequencing

Fecal sample collection, DNA extraction, PCR, and 16S rRNA library sequencing were performed as previously described (19). Fecal samples were collected prior to pellet implantation and once per week for the duration of the experiment. Fecal samples were frozen immediately after collection and stored at −80°C. DNA was extracted from the samples using the DNeasy PowerSoil Kit (Qiagen) according to the manufacturer’s protocol and stored at −80°C. PCR amplification was performed for the V4 hypervariable region of the 16S rRNA gene with primers 515F and 806R (35). The reverse primers contained unique 12-bp Golay barcodes that were incorporated into the PCR amplicons (35). Amplicon sequence libraries were prepared at the Scripps Research Institute Next Generation Sequencing Core Facility and sequenced on an Illumina MiSeq.

### Bioinformatics and Statistical Analysis of 16S rRNA gene sequences

Processing of sequences and OTU picking were performed using accessory scripts from QIIME version 1.9.1(add citation). Only forward reads were used from the Illumina sequencing data. Barcodes were extracted from the Illumina 16Ss fastq file using extract_barcodes.py and the data was demultiplexed and quality filtered using split_libraries_fastq.py with default parameters. OTU picking was performed using a *de novo* approach with the pick_de_novo_otus.py script, using Greengenes 13.8 as the reference database (add citation). The OTU table was then parsed using filter_otus_from_otu_table.py and any OTUs not present in at least 25% of samples were removed prior to downstream analysis.

### Fecal metabolite extraction and LC-MS/MS

Individual fecal samples were weighed to ensure they weighed at least 0.01 g per fecal sample. The fecal samples were transferred to 2 mL vial inserts to which a 1:10 volume of methanol (Optima LC/MS Grade Methanol, 67-56-1, Fisher Scientific) diluted in water 70:30 (Optima LC/MS Grade Water, 7732-18-5, Fisher Scientific) was added. The sample was then homogenized in a Qiagen TissueLyser and allowed to extract overnight at room temperature. After extraction, samples were briefly vortexed and incubated for 1 hour at room temperature before centrifugation at 10,000 g for 30s. Liquid chromatography was performed with ThermoScientific UltraMate 3000 Dionex. High-performance liquid chromatography (HPLC) was performed using a Phenomenex (Torrance, CA, USA) Luna 5 μm C18(2) HPLC column (2.0 mm × 250 mm) and ultra High-Performance Liquid Chromatography (UPLC) was performed using a Phenomenex Kinetex 2.6 μm C18 (30 × 2.10 mm) column with 20 μl of the extractions from the fecal pellets. A linear water–acetonitrile gradient (from 98:2 to 2:98 water:acetonitrile) containing 0.1% formic acid was used (HPLC: 54 min gradient; UPLC: 14 min gradient) with a flow rate of 0.2 ml min−1 for the HPLC analysis and 0.5 ml min−1 for the UPLC analysis. Tandem mass spectrometry (MS/MS) was performed using a Bruker Daltonics Maxis qTOF mass spectrometer equipped with a standard electrospray ionization source. Tuning of the mass spectrometer was done by infusion of Tuning Mix ES-TOF (Agilent Technologies) at a 3 μl min−1 flow rate. Lock mass internal calibration used a wick saturated with hexakis (1H,1H,3H-tetrafluoropropoxy) phosphazene ions (Synquest Laboratories, m/z 922.0098) located within the source for accuracy. The mass spectrometer was operated in data-dependent positive ion mode, automatically switching between full-scan MS and MS/MS acquisitions for both the HPLC and UPLC analyses. Full-scan MS spectra (m/z 50–2000) were acquired in the TOF and the top 10 most intense ions in a particular scan were fragmented using collision induced dissociation at 35 eV for +1 ion and 25 eV for +2 ions in the collision cell.

### Bioinformatics analysis of metabolites

Molecular networks were created by using the GNPS database online workflow at http://gnps.ucsd.edu, and the data set was used to search various MS/MS libraries available in the GNPS database by using the same workflow. The data set is available to the public at the online MassIVE repository of the GNPS database under MassIVE ID number MSV000081524. The molecular network used for analysis is available here: https://gnps.ucsd.edu/ProteoSAFe/status.jsp?task=d74cab73cc344821a59979f58269f3a8 Features were quantified using the mzMine-based feature finding algorithm with qTOF presets on the GNPS workflow. The features in the table were then filtered by removing features that were present in fewer than four samples, which removed approximately 50% of the total features. Metabolite annotations are based on MS/MS matches in the GNPS libraries and are therefore considered level two according to the metabolomics standards initiative (36).

### Shotgun metagenomic sequencing and preprocessing

800 ng of genomic DNA isolated from week 2 and week 5 samples was sonicated using an E220 focused ultrasonicator (Covaris) to produce 400 bp fragments which were purified using Agencourt AMPure XP beads (Beckman Coulter). A KAPA Hyper Prep Kit (Kapa Biosystems) was used to prepare Illumina libraries following the manufacturer’s instructions. Libraries were quality checked for their size and concentration with electrophoresis using a High Sensitivity D1000 kit on a 2200 TapeStation (Agilent). Prepared samples were sequenced by the Center for Advanced Technology at the University of California, San Francisco using an Illumina NovaSeq sequencer set to 150-bp paired-end reads. This produced an average of 109,084,767 reads per sample. Sequences from metagenomes were trimmed and filtered based on quality score, read length, and number of ambiguous nucleotides (N) using Fastp (version 0.19.6) (37). Adapters and PolyG tails (NovaSeq’s no signal indication) were automatically detected and removed. This preprocessing step removed < 1% of reads per sample, resulting in an average read length of 146 bp and a Q20 of 99.1%. Preprocessed paired-end reads were then mapped using Bowtie2 (version 2.2.6) to the mouse host genome (University of California, Santa Cruz *Mus musculus* genome; mm10) to remove host contamination (38). Mapped reads were removed with SAMtools (version 1.5) and unmapped reads were reconstructed to paired-end reads with BEDTools (version 2.25.0) (38, 39).

### Bioinformatics analysis of metagenome data

Taxonomic identification was performed using Centrifuge (version 1.0.3) against the NCBI non-redundant sequence database (40) and archaea, eukaryotes, and viral results were removed. Bacterial features (18,464) were passed through a filter that required species to be present in 90% of all samples and have an abundance of greater than .00001%, resulting in 3,539 bacterial species for analysis.

### Statistical analysis of metabolites, 16S rRNA genes and metagenomes

Analysis of the similarity among treatment groups was done via CAP using Bray-Curtis dissimilarity through the R package (version 1.26.1) (41). Differences among treatment groups was determined by permutational multivariate analysis of variance (PERMANOVA) using the python package Scikit-bio (version 0.5.5). Differential expressions of features were determined through the R package DESeq2 (version 1.18.1) using Wald’s test to find the log_2_ fold expression levels between treatment groups (42). Comparisons were made between P and LET treatments and LET and LET^ch^ treatments for each time point (week 2 and week 5).

### Multiomics analyses

To detect potential associations between the members of the gut microbial community and specific metabolites, two multiomics feature tables were created: 1) Metabolomes combined with 16S rRNA sequencing and 2) Metabolomes and metagenomes. Multiomics data were analyzed via CAP based on Bray-Curtis dissimilarities through Phyloseq (version 1.26.1) (41). BAs and bacterial species with the largest log_2_ fold magnitude as indicated by DESeq2 were used as features in a random forest supervised learning model analysis via Scikit-learn (version 0.20.1) in Python (42, 43). All models were optimized to have the lowest out-of-bag error. Following the same comparison structures in the DESeq2 analysis, the top 10 features with the highest Gini importance index for all comparisons were selected for a correlation analysis. To combine multiomics datasets at very different scales for correlation analysis, we transformed the datasets using the centered log-ratio (clr) approach. Zero-replacement was performed with pseudo-counts method from the R package zCompositions version 1.3.3. Spearman-rank correlations analysis was performed in Python pandas (version 0.25.1) with clr-transformed BA counts and clr-transformed bacterial species counts from the metagenomic data for P, Let, and LET^ch^ at weeks 2 and 5. Heatmaps were generated using the Python seaborn package (version 0.9.0) (44, 45).

### Data availability

16S rRNA gene sequences used in this study are available via the European Nucleotide Archive (Study Accession Number PRJEB29583), Shotgun Metagenomes are available via the European Nucleotide Archive (Study Accession Number PRJEB40312; Metabolomics data is available online MassIVE repository of the GNPS database under MassIVE ID number MSV000081524.

## SUPPLEMENTAL MATERIAL

The metadata and all code used to analyze and visualize data are available at: https://github.com/bryansho/PCOS_WGS_16S_metabolome.

## ACKNOWLEDGEMENTS

We thank members of the Thackray and Kelley labs for insightful comments and suggestions. We also thank Pieter Dorrestein for providing training and access to the facilities at the mass spectrometry center. Gut microbial metagenomes were sequenced by the University of California, San Diego Institute for Genomic Medicine (P30DK063491).

This work was funded by the National Institute of Child Health and Human Development through a cooperative agreement as part of the National Centers for Translational Research in Reproduction and Infertility (Grant P50 HD012303, to V.G.T.). V.G.T. and S.T.K. were also funded by Grant R01 HD095412. P.J.T. was funded by the University of California, San Diego Microbial Sciences Research Initiative Fellowship and the San Diego State University Achievement Rewards for College Scientists Foundation.

## REFERENCES

1. Lizneva D, Suturina L, Walker W, Brakta S, Gavrilova-Jordan L, Azziz R. 2016. Criteria, prevalence, and phenotypes of polycystic ovary syndrome. Fertil Steril.

2. Palomba S, De Wilde MA, Falbo A, Koster MPH, La Sala GB, Fauser BCJM. 2015. Pregnancy complications in women with polycystic ovary syndrome. Hum Reprod Update.

3. Fauser BCJM. 2004. Revised 2003 consensus on diagnostic criteria and long-term health risks related to polycystic ovary syndrome. Fertil Steril.

4. Vink JM, Sadrzadeh S, Lambalk CB, Boomsma DI. 2006. Heritability of polycystic ovary syndrome in a Dutch twin-family study. J Clin Endocrinol Metab.

5. De Leo V, Musacchio MC, Cappelli V, Massaro MG, Morgante G, Petraglia F. 2016. Genetic, hormonal and metabolic aspects of PCOS: An update. Reprod Biol Endocrinol.

6. Franks S, McCarthy MI, Hardy K, Skakkebæk NE, Aitken RJ, Swan S, De Muinck Keizer-Schrama S. 2006. Development of polycystic ovary syndrome: Involvement of genetic and environmental factorsInternational Journal of Andrology.

7. Anderson AD, Solorzano CMB, McCartney CR. 2014. Childhood obesity and its impact on the development of adolescent PCOS. Semin Reprod Med.

8. Azziz R, Carmina E, Chen Z, Dunaif A, Laven JSE, Legro RS, Lizneva D, Natterson-Horowtiz B, Teede HJ, Yildiz BO. 2016. Polycystic ovary syndrome. Nat Rev Dis Prim.

9. Moran LJ, Misso ML, Wild RA, Norman RJ. 2010. Impaired glucose tolerance, type 2 diabetes and metabolic syndrome in polycystic ovary syndrome: A systematic review and meta-analysis. Hum Reprod Update.

10. Barber TM, Wass JAH, McCarthy MI, Franks S. 2007. Metabolic characteristics of women with polycystic ovaries and oligo-amenorrhoea but normal androgen levels: Implications for the management of polycystic ovary syndrome. Clin Endocrinol (Oxf).

11. Moghetti P, Tosi F, Bonin C, Di Sarra D, Fiers T, Kaufman JM, Giagulli VA, Signori C, Zambotti F, Dall’Alda M, Spiazzi G, Zanolin ME, Bonora E. 2013. Divergences in insulin resistance between the different phenotypes of the polycystic ovary syndrome. J Clin Endocrinol Metab.

12. Lindheim L, Bashir M, Münzker J, Trummer C, Zachhuber V, Leber B, Horvath A, Pieber TR, Gorkiewicz G, Stadlbauer V, Obermayer-Pietsch B. 2017. Alterations in gut microbiome composition and barrier function are associated with reproductive and metabolic defects in women with polycystic ovary syndrome (PCOS): A pilot study. PLoS One.

13. Liu R, Zhang C, Shi Y, Zhang F, Li L, Wang X, Ling Y, Fu H, Dong W, Shen J, Reeves A, Greenberg AS, Zhao L, Peng Y, Ding X. 2017. Dysbiosis of gut microbiota associated with clinical parameters in polycystic ovary syndrome. Front Microbiol.

14. Valdes AM, Walter J, Segal E, Spector TD. 2018. Role of the gut microbiota in nutrition and health. BMJ.

15. Durack J, Lynch S V. 2019. The gut microbiome: Relationships with disease and opportunities for therapy. J Exp Med.

16. Torres PJ, Siakowska M, Banaszewska B, Pawelczyk L, Duleba AJ, Kelley ST, Thackray VG. 2018. Gut Microbial Diversity in Women with Polycystic Ovary Syndrome Correlates with Hyperandrogenism. J Clin Endocrinol Metab.

17. Kauffman AS, Thackray VG, Ryan GE, Tolson KP, Glidewell-Kenney CA, Semaan SJ, Poling MC, Iwata N, Breen KM, Duleba AJ, Stener-Victorin E, Shimasaki S, Webster NJ, Mellon PL. 2015. A Novel Letrozole Model Recapitulates Both the Reproductive and Metabolic Phenotypes of Polycystic Ovary Syndrome in Female Mice1. Biol Reprod.

18. Kelley ST, Skarra D V., Rivera AJ, Thackray VG. 2016. The gut microbiome is altered in a Letrozole-Induced mouse model of polycystic ovary syndrome. PLoS One.

19. Torres PJ, Ho BS, Arroyo P, Sau L, Chen A, Kelley ST, Thackray VG. 2019. Exposure to a Healthy Gut Microbiome Protects Against Reproductive and Metabolic Dysregulation in a PCOS Mouse Model. Endocrinology2019/03/30. 160:1193–1204.

20. Arroyo P, Ho BS, Sau L, Kelley ST, Thackray VG. 2019. Letrozole treatment of pubertal female mice results in activational effects on reproduction, metabolism and the gut microbiome. PLoS One2019/10/01. 14:e0223274.

21. Guo Y, Qi Y, Yang X, Zhao L, Wen S, Liu Y, Tang L. 2016. Association between polycystic ovary syndrome and gut microbiota. PLoS One.

22. Jahan S, Abid A, Khalid S, Afsar T, Qurat-Ul-Ain, Shaheen G, Almajwal A, Razak S. 2018. Therapeutic potentials of Quercetin in management of polycystic ovarian syndrome using Letrozole induced rat model: A histological and a biochemical study. J Ovarian Res.

23. Walter J, Armet AM, Finlay BB, Shanahan F. 2020. Establishing or Exaggerating Causality for the Gut Microbiome: Lessons from Human Microbiota-Associated Rodents. Cell.

24. Ridaura VK, Faith JJ, Rey FE, Cheng J, Duncan AE, Kau AL, Griffin NW, Lombard V, Henrissat B, Bain JR, Muehlbauer MJ, Ilkayeva O, Semenkovich CF, Funai K, Hayashi DK, Lyle BJ, Martini MC, Ursell LK, Clemente JC, Van Treuren W, Walters WA, Knight R, Newgard CB, Heath AC, Gordon JI. 2013. Gut microbiota from twins discordant for obesity modulate metabolism in mice. Science (80-).

25. Qi X, Yun C, Sun L, Xia J, Wu Q, Wang Y, Wang L, Zhang Y, Liang X, Wang L, Gonzalez FJ, Patterson AD, Liu H, Mu L, Zhou Z, Zhao Y, Li R, Liu P, Zhong C, Pang Y, Jiang C, Qiao J. 2019. Gut microbiota–bile acid–interleukin-22 axis orchestrates polycystic ovary syndrome. Nat Med.

26. Caruso R, Ono M, Bunker ME, Núñez G, Inohara N. 2019. Dynamic and Asymmetric Changes of the Microbial Communities after Cohousing in Laboratory Mice. Cell Rep.

27. Wang M, Zhang Y, Miller D, Rehman NO, Cheng X, Yeo JY, Joe B, Hill JW. 2020. Microbial Reconstitution Reverses Early Female Puberty Induced by Maternal High-fat Diet During Lactation. Endocrinology.

28. Wahlström A, Sayin SI, Marschall HU, Bäckhed F. 2016. Intestinal Crosstalk between Bile Acids and Microbiota and Its Impact on Host Metabolism. Cell Metab. Cell Press.

29. Ruiz L, Margolles A, Sánchez B. 2013. Bile resistance mechanisms in Lactobacillus and Bifidobacterium. Front Microbiol 4:396.

30. Urdaneta V, Casadesús J. 2017. Interactions between bacteria and bile salts in the gastrointestinal and hepatobiliary tracts. Front Med. Frontiers Media S.A.

31. Tian Y, Gui W, Koo I, Smith PB, Allman EL, Nichols RG, Rimal B, Cai J, Liu Q, Patterson AD. 2020. The microbiome modulating activity of bile acids. Gut Microbes 11:979–996.

32. van Best N, Rolle-Kampczyk U, Schaap FG, Basic M, Olde Damink SWM, Bleich A, Savelkoul PHM, von Bergen M, Penders J, Hornef MW. 2020. Bile acids drive the newborn’s gut microbiota maturation. Nat Commun.

33. O’Flaherty S, Briner Crawley A, Theriot CM, Barrangou R. 2018. The Lactobacillus Bile Salt Hydrolase Repertoire Reveals Niche-Specific Adaptation. mSphere.

34. Wu TR, Lin CS, Chang CJ, Lin TL, Martel J, Ko YF, Ojcius DM, Lu CC, Young JD, Lai HC. 2019. Gut commensal Parabacteroides goldsteinii plays a predominant role in the anti-obesity effects of polysaccharides isolated from Hirsutella sinensis. Gut.

35. Caporaso JG, Lauber CL, Walters WA, Berg-Lyons D, Huntley J, Fierer N, Owens SM, Betley J, Fraser L, Bauer M, Gormley N, Gilbert JA, Smith G, Knight R. 2012. Ultra-high-throughput microbial community analysis on the Illumina HiSeq and MiSeq platforms. ISME J.

36. Sumner LW, Amberg A, Barrett D, Beale MH, Beger R, Daykin CA, Fan TW-M, Fiehn O, Goodacre R, Griffin JL, Hankemeier T, Hardy N, Harnly J, Higashi R, Kopka J, Lane AN, Lindon JC, Marriott P, Nicholls AW, Reily MD, Thaden JJ, Viant MR. 2007. Proposed minimum reporting standards for chemical analysis. Metabolomics.

37. Chen S, Zhou Y, Chen Y, Gu J. 2018. Fastp: An ultra-fast all-in-one FASTQ preprocessorBioinformatics.

38. Langmead B, Salzberg S. 2013. Bowtie2. Nat Methods.

39. Quinlan AR, Hall IM. 2010. BEDTools: A flexible suite of utilities for comparing genomic features. Bioinformatics.

40. Kim D, Song L, Breitwieser FP, Salzberg SL. 2016. Centrifuge: Rapid and sensitive classification of metagenomic sequences. Genome Res.

41. McMurdie PJ, Holmes S. 2013. Phyloseq: An R Package for Reproducible Interactive Analysis and Graphics of Microbiome Census Data. PLoS One.

42. Love MI, Huber W, Anders S. 2014. Moderated estimation of fold change and dispersion for RNA-seq data with DESeq2. Genome Biol.

43. Pedregosa F, Varoquaux G, Gramfort A, Michel V, Thirion B, Grisel O, Blondel M, Prettenhofer P, Weiss R, Dubourg V, Vanderplas J, Passos A, Cournapeau D, Brucher M, Perrot M, Duchesnay É. 2011. Scikit-learn: Machine learning in Python. J Mach Learn Res.

44. McKinney W. 2011. pandas: a Foundational Python Library for Data Analysis and Statistics. Python High Perform Sci Comput.

45. Waskom M. 2018. Seaborn: Statistical Data Visualization — Seaborn 0.9.0 Documentation. Sphinx 174.

